# Active Sampling in Primate Vocal Interactions

**DOI:** 10.1101/2023.12.05.570161

**Authors:** Thiago T. Varella, Daniel Y. Takahashi, Asif A. Ghazanfar

## Abstract

Active sensing is a behavioral strategy for exploring the environment. In this study, we show that contact vocal behaviors can be an active sensing mechanism that uses sampling to gain information about the social environment, in particular, the vocal behavior of others. With a focus on the real-time vocal interactions of marmoset monkeys, we contrast active sampling to a vocal accommodation framework in which vocalizations are adjusted simply to maximize responses. We conducted simulations of a vocal accommodation and an active sampling policy and compared them with real vocal exchange data. Our findings support active sampling as the best model for marmoset monkey vocal exchanges. In some cases, the active sampling model was even able to predict the distribution of vocal durations for individuals. These results suggest a new function for primate vocal interactions in which they are used by animals to seek information from social environments.

**Significance:** We found that marmoset monkeys use vocal exchanges with conspecifics in the same way that bats and electric fish use their self-generated signals: to sample the information space. The difference is that, while bats, electric fish and other echolocating animals emit signals and then interpret that signal’s reflection, for vocalizing marmoset monkeys the “reflection” is the vocal response from conspecifics. We show that marmosets produce variable vocalizations as a means to sample the acoustic space in which they can elicit a conspecific response. They are actively sampling--learning about their dyadic partner like a bat learns about its prey.

## Introduction

Contact calls are produced by a variety of animals, particularly when they are out of sight of one another. The functions of contact calling seem to be context dependent. Among primate species, the same vocalizations are variously used during territorial encounters, mate attraction and isolation (1). In all contexts, contact calls produced by one individual typically elicit a similar vocalization by another. There are two common assumptions as to the immediate purpose of these vocal exchanges. The first is that the goal of each individual is to maximize the probability of a response from the other (e.g., frogs: (2, 3); monkeys: (4, 5)). The second assumption is that, in some cases, vocal plasticity is used to either adjust these vocalizations acoustically to signal social closeness or to optimize signal transmission in noisy environments. This, again, has the purpose of increasing the probability of a vocal response (a phenomenon known as *vocal accommodation* (6)). Here, we consider another possibility: vocal exchanges are a form of active sensing used to gain information about conspecifics (that is, they are *active sampling*) (7).

Active sensing is the use of motor output--like moving your eyes across a scene or palpating an object in the dark--to collect sensory signals (8). In the domain of acoustics, echolocation is the paradigmatic case of active sensing as stimulus energy (a vocalization) is generated by the subject to detect, localize, and discriminate objects in the environment via the energy’s reflection. Sound-based active sensing by bats and other echolocating animals is a highly specialized form of communication, whereby an animal serves as both the sender and receiver of sensory energy. For example, bats emit one type of vocal signal to search for prey and, when a promising echo signal is returned, they change their vocal output to better sample what it might be (9, 10). We propose here that contact calling vocal behavior is akin to this process. Like echolocating in the dark, contact calls are in essence signals generated to find something (in this case, conspecifics). However, instead of the vocal sound generating an echo reflected off prey, these vocalizations may elicit a vocal signal from a conspecific. Like echolocation signals used for prey capture, contact calls are used to detect, localize and discriminate conspecifics (“is anyone out there?”, “who are you?” and “where are you?”), *i*.*e*. to learn more about the social environment (11).

Marmoset monkeys are one model species where there has been much focus on the use of their contact calls in vocal exchanges (Figure 1). When by themselves, adult marmosets will produce contact calls and will do so approximately every 10 seconds as a function of an autonomic nervous system rhythm (12, 13). When auditory contact is made, two marmosets will communicate with multiple back-and-forth exchanges of contact calls (13). When the distance between them changes or if they become visible to each other, then marmosets adjust the latency, loudness, and/or duration of their contact calls and/or switch to producing different affiliative call types (14, 15). These and other data demonstrate that marmoset vocal production remains flexible or *plastic* throughout adulthood (16-18). Here, we test whether the real-time contact calling behavior of marmoset monkeys is consistent with active sampling. Are they interrogating their social environment using a “question-and-answer” strategy?

**Figure 1.**
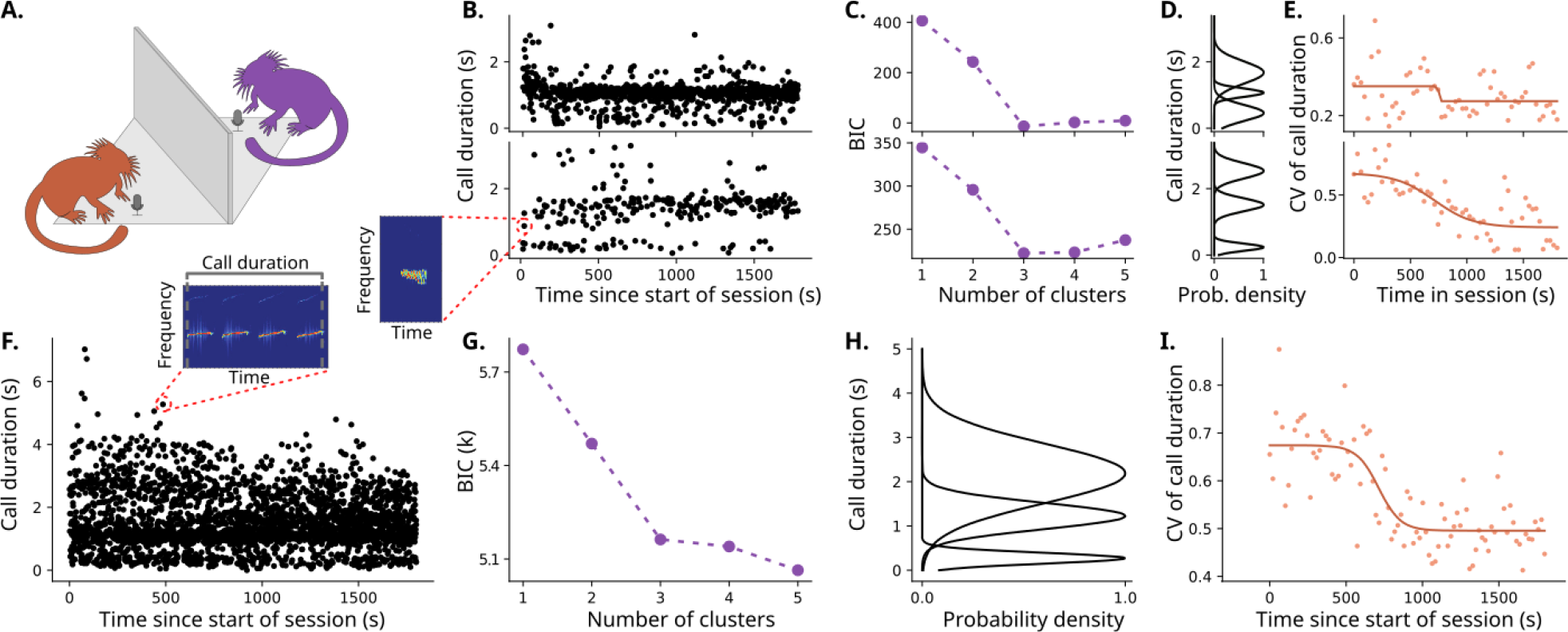
Vocal behavior within a single dyadic interaction is diverse and dynamic. **A**. Experimental set-up of the occluded and dyadic vocal interactions. **(B-E)**. Vocal properties for two individuals, individual A at the top and individual B at the bottom. **B**. Scatter plot for each individual with the time that each call was produced in the x-axis and the call duration of said call in the y-axis. The call duration is calculated as the total duration of a sequence of vocal syllables (continuous bouts of vocalization) with less than 1 second between each other (13), as exemplified in the spectrogram. **C**. Value of Bayesian Information Criterion (BIC) versus the number of components for each individual when clustering with a Gaussian mixture model the duration of all vocalizations produced after ¼ of the session. For both individuals, 3 is the optimal number of clusters. **D**. Probability density function of each of the clusters for each individual, as determined by the Gaussian mixture model of the optimal number of clusters. **E**. Dynamics of the time-binned coefficient of variation (CV, standard deviation divided by the mean) of the call durations for each individual, calculated via splitting the data into time windows, and calculating the CV in each window. The solid line represents a sigmoid fit of the obtained CVs. **F**. Scatter plot of vocal durations for the population of 6 individuals throughout the sessions. **G**. BIC versus the number of clusters showing that 3 is also the optimal number of clusters of call durations considering the population as a whole. **H**. Probability density function illustrating what are the clusters. **I**. Dynamics of CV of call duration for the population. Solid line is a sigmoid fit.

## Results

Active sampling involves gathering information relevant for a task (7). One proposal for active sampling suggests that animals in complex environments with incomplete information use a belief-based policy (19), where “belief” here means the state of an animal’s knowledge at that moment. In a belief-based system, information has value, and this value can motivate exploration under conditions in which they must consider many alternatives and where the potential rewards are not known beforehand. In the context of contact calling by marmoset monkeys who, under natural conditions, live in dense, tropical rainforests where they cannot easily see conspecifics, we hypothesize that vocalizing is a way of seeking information about conspecifics.

It is well known that in echolocation by bats, they use their vocalization as a way of seeking information about potential food in their surroundings. Importantly, once food has been detected, the duration of their vocalizations changes as function of various stages of the approach and acquisition of the food (e.g., an insect or another source of energy). Bats start by emitting longer vocalizations called search or orienting calls, which is followed by shorter approach calls as they get closer to the food, ending the process with very short duration terminal calls. This shows that call duration is an important acoustic feature for active sensing in bats, demonstrating that bats produce different vocalizations in the single context of foraging for food (20, 21). To assess the hypothesis that a similar phenomenon happens with marmosets, we measured and analyzed different call durations in the single context of vocal exchanges with an out of sight conspecific. We used raw recording data collected from an earlier study (15); the vocalizations were from six adult marmoset monkeys, split into three pairs. The pair was put into a sound-attenuated testing room where one marmoset was placed in one corner and another in the other corner, separated by an acoustically transparent visual occluder (Figure 1A). This context reliably elicits vocal exchanges between marmosets (13, 15). As in previous studies, vocal responses were defined as a vocalization from a different marmoset that occurred within 12 seconds of the start of the first vocalization (13, 15).

When we plot the duration of the vocalizations emitted over time for individuals (Figure 1B) or the population (Figure 1F), we can see that, qualitatively, most of the calls with a longer duration are emitted in the beginning of the session. This is similar to bats that reduce their call duration as they approach their targets. We quantified this using Bayesian Information Criteria and a Gaussian Mixture Model (Figure 1C and D for the individuals and Figure 1G and H for the population).

These data show that distinct groups of vocal durations are evident. We know that different vocal durations represent different call types (22) and these vocalizations are all affiliative in this context (15). Why are marmosets producing different call types in the same context? Vocalizations are shaped by energetic and biomechanical constraints (23). Thus, an observer might expect that, if the marmoset knows that the receiver is present after hearing them vocally respond and they begin responding at an optimal rate, then there will come a time when there is no need to emit different call types. The idea that the marmoset may be acquiring such information throughout the session is supported by the observation that the coefficient of variation of call durations is reduced over time (Figures 1E and I). This pattern is highly reminiscent of active sampling by bats described above. Thus, we now try to provide a quantitative model for what this would look like for marmoset monkeys.

To investigate this question, we began by simulating two scenarios for how a pair of agents would change the duration of their vocalizations during one session of vocal exchanges. Figure 2 illustrates these putative processes using two different behavioral policies. In the first scenario--vocal accommodation (Figure 2A)--the agents change their vocalizations so that they produce calls with durations that maximize the perceived likelihood of getting a response. For example, if they vocalize and get no response, they might produce a longer duration vocalization, making it more likely that the other agent hears it. If the agent successfully gets a response after a longer call, then they would produce a call with similar duration next time. This would increase the probability of response. In this scenario, the agents can learn from their previous vocalizations produced during the session.

**Figure 2.**
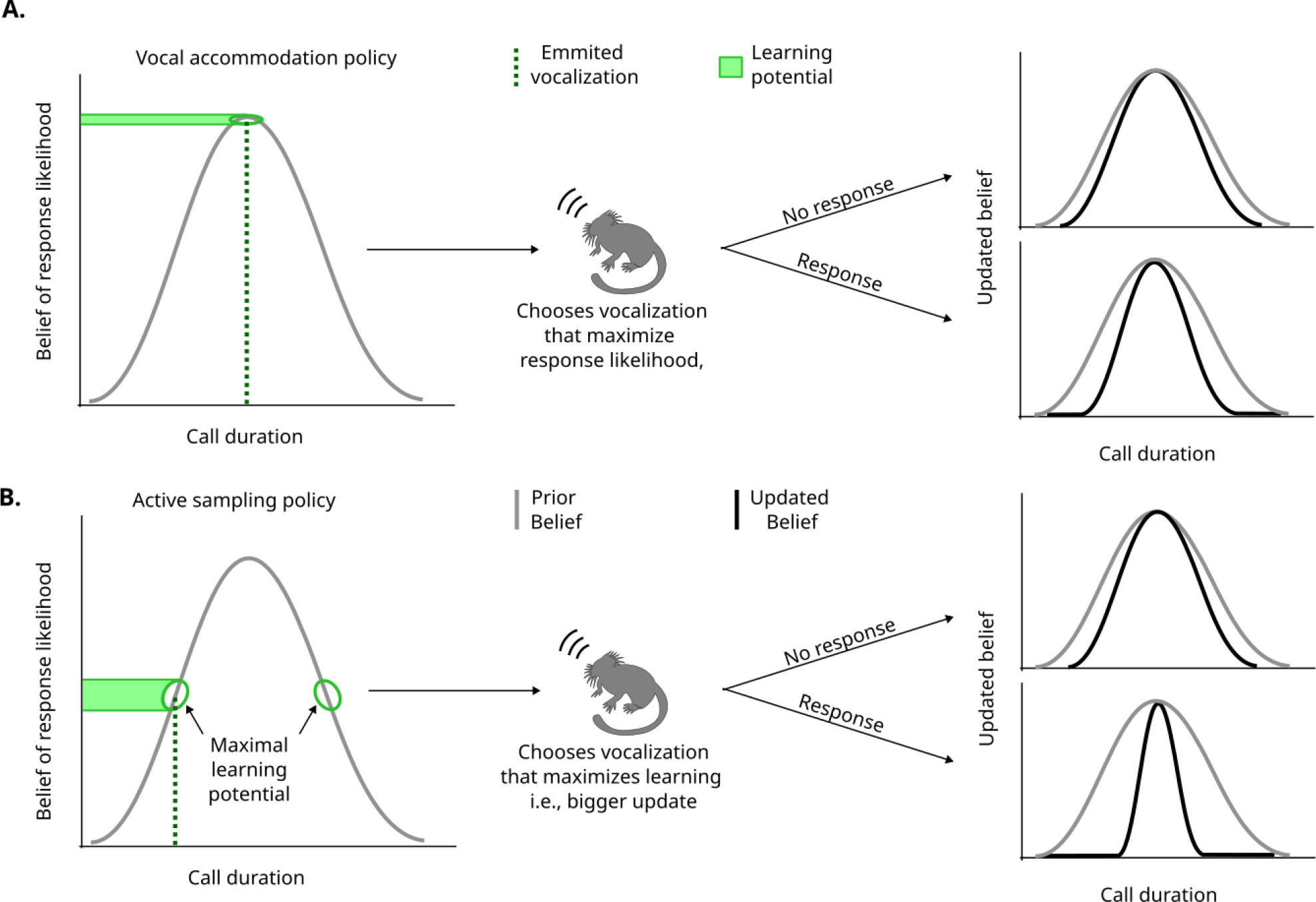
Different vocalizations chosen by a policy lead to different degrees of information acquisition. In both models, the agent starts with a belief (represented by a distribution) of what call duration leads to the highest response probability. Using a policy applied to this belief (i.e. the rules determining what action to take), the agent emits a vocalization with a specific duration. Based on the information it gets from hearing or not a response, it updates the belief using the Bayes rule. **A**. In the vocal accommodation policy, the vocalization is chosen to maximize the immediate response probability (i.e., the duration corresponding to the peak of the belief). **B**. Another possibility for the policy is the active sampling policy, in which the vocalization is chosen to maximize the learning (see methods). In the example in the figure, the policy leads to vocalizations on the highest slopes of the belief, rather than the peak, because they have a higher learning potential, achieving an update in the belief with higher magnitude and thus a narrower updated belief than the vocal accomodation model.

The distribution on the left of Figure 2A and 2B represents a “belief” acquired prior to (or during) the session, such as over the course of development of the agent (24, 25). This belief informs the agent how likely it is to get a response depending on certain acoustic features (call duration, in this case) of their vocalization. The vocalization chosen for this policy is at the peak of the belief distribution, meaning they will choose the vocalization that they think is the most likely to get a response. Notice, however, that by choosing the vocalization in the peak of the belief distribution, small changes on the choice of the vocalization (x-axis) lead to small changes in the response rate (y-axis). This is because of the flatness of the response rate around the peak. This is consistent with the notion that the agent is more “confident” about what the response rate should be in this area, and, therefore, there is a narrow learning potential.

If the belief correctly describes the response likelihood, it is more likely to get a response when vocalizing in the peak, but if the belief is incorrect, vocalizing in the peak could lead to suboptimal calling behavior and a reduced chance to improve. A more robust strategy for information seeking would be to choose a vocalization with a duration that has more potential for learning something new. In the second scenario—the active sampling policy (Figure 2B)--the agents change their vocal behavior so that they are more likely to learn something new. For example, even if specific vocalizations are already leading to consistent responses, the agent will try to vocalize longer or shorter as a way of learning something new that could lead to, perhaps, a higher response likelihood later on. Given the distribution of the belief, the agent will not necessarily select the vocalization in the peak (*i*.*e*., the one with highest perceived likelihood to get a response). One example of a vocalization that could be chosen in this scenario is indicated by the dotted line in Figure 2B. In this area, small variations in the chosen vocalization (perturbations on the x-axis of the distribution) lead to big changes in the expected response rate (green area by the y-axis). The height of the green area can serve as an intuition to the agent’s learning potential, since exposure to different response rates might help the agent better represent the correct distribution. For this specific distribution, this is equivalent to the area with the highest slope. A consequence of choosing the vocalization with highest learning potential is that the update in the belief will be larger, as observed in the updated belief distributions on the right. Here, when the agent gets a response, the updated distribution is a lot narrower, representing a big increase in confidence. Because of the symmetry of this belief distribution, there are two regions with maximal learning potential.

We conducted simulations of affiliative vocal exchanges between agent dyads (Figure 1A).

The simulations used Bayesian inference to investigate the two alternative behavioral policies described above. Specifically, we applied the Bayes rule to the belief of the optimal vocalization given a newly sampled vocalization and given a variable that represents the presence or absence of a response. Both agents in the simulation use the same policy (Methods). The policies relate to two different ways of thinking about why marmosets produce the vocalizations they do during vocal exchanges with a conspecific. The first policy considers that they are trying to maximize the probability of a response from their communication partner based on their prior knowledge. This policy is consistent with matching their vocalizations to their partner’s vocalizations. While marmosets do exhibit vocal accommodation like this over the course of weeks (and as a function of vocalization type) (18), it is unknown to what degree they do so on shorter timescales.

In Figure 3, each row is the result of implementing one of the two policies: vocal accommodation and active sampling. Figure 3A-C shows that the vocal accommodation policy converges mainly to one big peak along with a second, broader peak, while the active sampling policy (Figure 3E-G) converges from a broad distribution to two main narrow peaks and a third broader peak. The two main peaks in the active sampling policy are the result of the agents trying to explore an area around the vocalization with highest learning potential (Figure 2B). The vocal accommodation policy leads to only one sharp peak since the vocalization converges to the perceived optimal duration. In both cases, the broader peak is related to vocalizations that are important for the initial wider exploration of the different call durations (*i*.*e*., information search). The elbow method of cluster analysis was done in Figures 3B and F to algorithmically determine the number of clusters. Figures 3D and 3H show that both the vocal accommodation and the active sampling policies exhibit a transition in how variable the vocalization is throughout a session, although the transition in the active sampling policy seems sharper.

**Figure 3.**
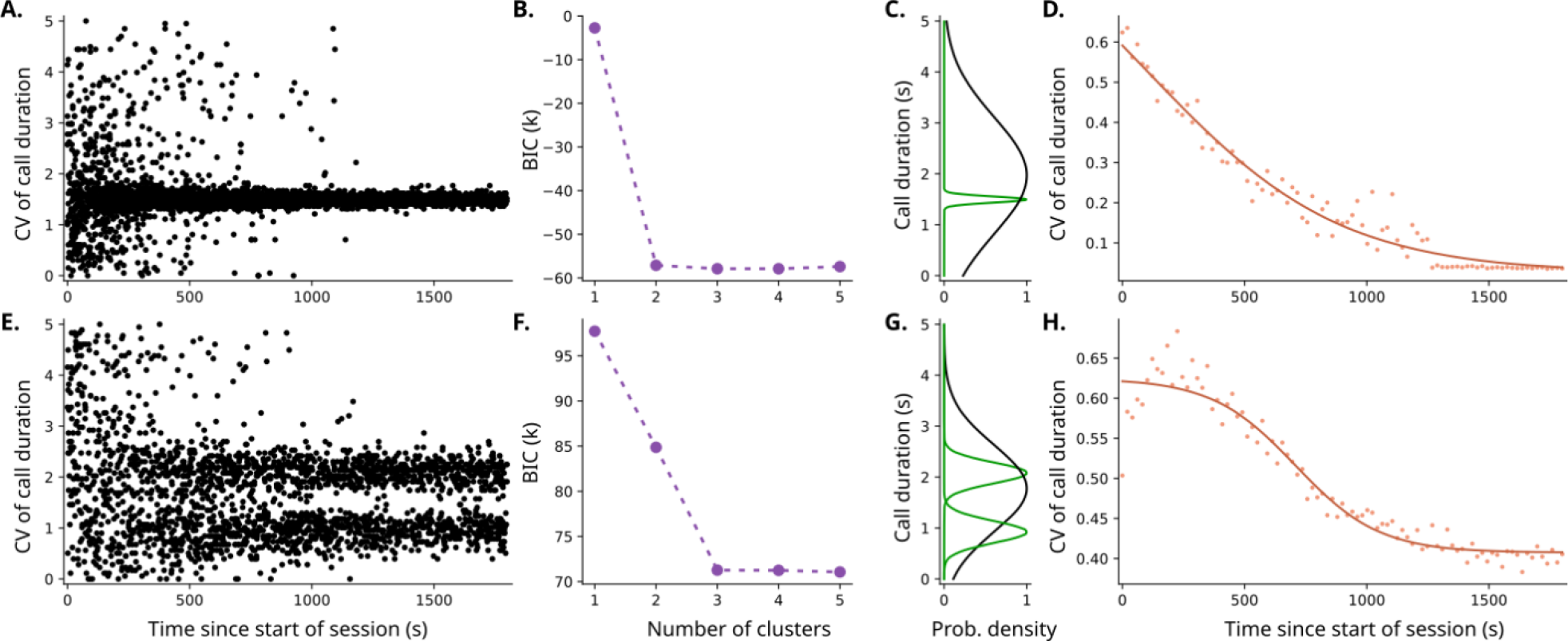
The active sampling model predicts diversity (3 clusters) and dynamics (sudden transition) of vocal interaction. **A**. Scatter plot of the simulated call durations in an interaction between two agents using a vocal accommodation policy. **B**. Value of Bayesian information criterion (BIC) versus the number of components when Gaussian mixed models are used to cluster the call durations after ¼ of the sessions simulated by the vocal accommodation policy. The optimal number of clusters defined via elbow method is 2. **C**. Probability density function of each of the clusters from the vocal accommodation model, using the optimal number of clusters. **D**. Dynamics of the time-binned coefficient of variation (CV, standard deviation divided by the mean) of the vocal accommodation simulation, calculated via splitting the data into bins in time, and calculating the CV in each bin. The solid line represents a sigmoid fit of the CVs. **E**. Scatterplot of the simulated call durations in an interaction between two agents using the active sampling policy. **F**. BIC vs number of components when clustering vocal durations with a Gaussian mixture model from calls taken after ¼ of the sessions in active sampling simulation. **G**. Probability density function of each of the clusters from the active sampling model, using the optimal number of clusters given by the elbow method. **H**. Dynamics of the time-binned CV of the active sampling simulation, along with a sigmoid fit.

Next, we compared what we observed in these simulations with the real marmoset vocal exchanges in Figure 1. Figures 1G and 1H show the same number of clusters as in Figures 3F and 3G (for the active sampling policy), while in Figure 1I we observe a sharp transition similar to the one observed in Figure 3H for active sampling. This suggests that the active sampling is the best model to concomitantly describe both the dynamics and the diversity of vocal durations when compared to the vocal accommodation policy. Both policies include the idea that the marmoset is updating their belief of the best vocalization in a given context, but the active sampling policy would mean that this update is not just passive, but that information is being actively pursued by the marmoset.

As predicted by the idea that the marmosets are actively sampling throughout a vocal interaction session, Figure 4A shows a statistically significant increase in the proportion of calls that receive a response from their partners as the session progresses (bootstrapped linear fit had positive slope with p = 0.042). There is a particularly sharp increase in the beginning of the curve, similarly to what we observe from the active sampling simulation. If the active sampling policy is used, there should be two clusters of call durations around the peak of an individual belief. We can plot the experience that a marmoset has with the proportion of responses heard given different call durations emitted, which is similar to what the belief should represent. The plots in Figures 4C and 4E show such curves for marmosets A and B (the same individuals from Figure 1), and they are similar to the beliefs illustrated in Figure 2. As expected, Figures 4B and 4D show the clusters of vocal durations emitted by the same individuals, and the peaks of the clusters bracket the proportion of vocalizations that elicit a response.

**Figure 4.**
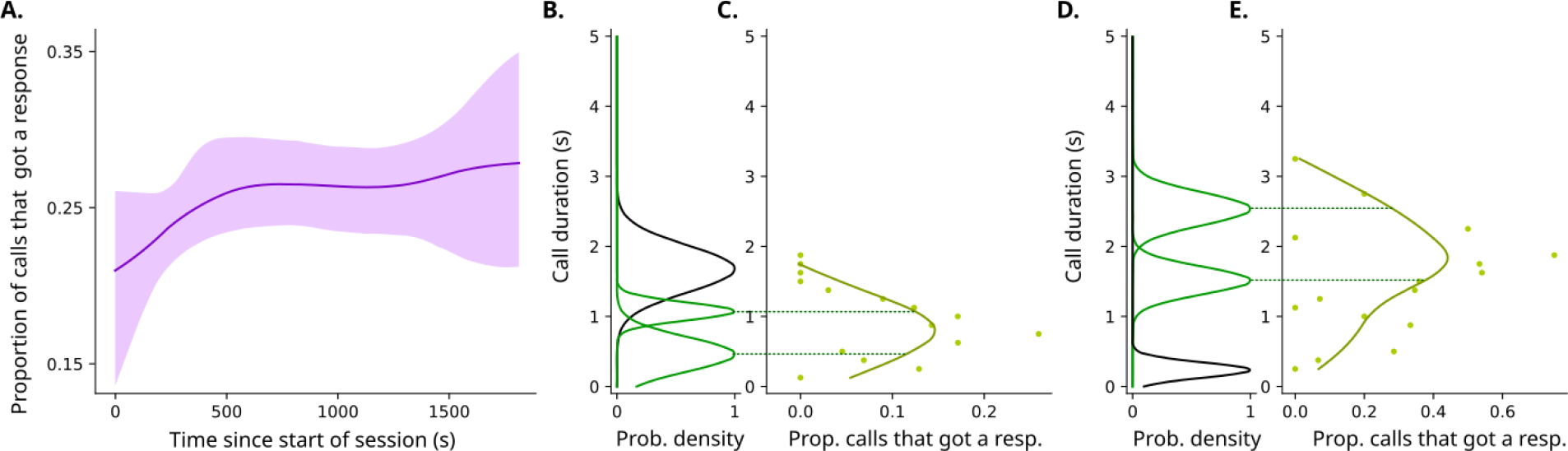
Marmoset behavior matches predictions from active sampling model. **A**. Graph showing increase in proportion of calls that get a response as the session progresses for the population of 6 individuals. **B**. Probability density function of each of the clusters individual A, as determined by the Gaussian mixture model of the optimal number of clusters shown in Fig. 1D. **C**. Proportion of calls from individual A that got a response (in the x-axis) given different call durations in the y-axis. The dotted line shows alignment of the vocal clusters similarly to the curve, as predicted by the active sampling model in Fig. 2B. **D**. Probability density function of each of the clusters individual B, as determined by the Gaussian mixture model of the optimal number of clusters shown in Fig. 1D. **E**. Proportion of calls from individual B that got a response (in the x-axis) given different call durations in the y-axis. Dotted line illustrates alignment of the vocal clusters with the curve.

## Discussion

It is impossible, and would be highly inefficient, to build a complete representation of all the sensory information in a particular context. Thus, sampling is a necessity for organisms that can sense much more information than they can fully process. It is a means to acquire in small amounts the very rich information coming in from the different sensory systems (7). Active sampling is observed across different sensory modalities, such as smelling, active vision, and active touch (26-28). The idea that these senses are used actively by most animals is ubiquitous, but hearing is often left behind in these discussions (29). Considerations of actively seeking information using hearing and vocal behavior is typically confined to echolocating animals. Our results showed that, in real time, marmosets out of sight of one another primarily use active sampling to guide their vocal production. This suggests that contact vocalizations can be used as an active sensing mechanism similar (though not identical) to echolocation in other animals.

It is important to note that the active sampling policy is not mutually exclusive of vocal accommodation in a broader, long-term sense. Our paper only analyzed short-term vocal changes (within the range of 30 minutes). The differential with active sensing is that it allows for more flexibility in a rapidly changing environment. As it oscillates around the optimal vocalization, when such optimal vocalization changes (e.g., in the presence of another animal, or when the marmoset vocalizing changes position), the agent is already primed to probe the areas around what it believed to be optimal, instead of insisting on the same vocalization. In other words, it can lead to quicker accommodation, when the environment changes. This explains why active sensing is observed in short term interactions even though accommodation is still observed in the span of days and months in marmoset monkeys (17, 18).

Marmoset vocal behavior can be characterized as a balance between energy expenditure and information gain (30). Such energy-information balancing is common among other established models of active sampling. For example, the amount of energy a weakly electric fish expends in movement is a function of how much sensory information it seeks to gain via electrolocation (31). Weakly electric fishes also make communication decisions that are a necessary part of the active sampling mechanism—a means to continually probe the environment (32). Marmoset monkeys also have a diverse vocal repertoire whose function is not completely understood. For example, they use multiple types of affiliative vocalizations that are seemingly produced according to physical distance from conspecifics (15, 33, 34), but sometimes produce (sub-optimally, in terms of energy usage) loud and long duration affiliative vocalizations in close contexts. We believe that, like electric fish, marmosets are using their contact vocalizations to probe the social environment. The information they are acquiring could be related to individual identity, indexical cues such as age, gender and body size, and/or the location of the individual: all such information is known to be present in the contact vocalizations of marmoset monkeys (35-37).

Our findings represent a departure from previous research on contact vocalizations not only for the claim that we believe they are being used for active sampling but also because we are suggesting that in real time, vocalizations are not emitted solely to maximize the perceived probability of a conspecific’s response. The assumption that a maximal response rate is a goal of the contact communication system is embedded in previous work which includes the literature on vocal accommodation (6), turn-taking (including our previous studies (25)), and general observations about vocal behavior (3). In the simulation of the accommodation policy, vocalizations quickly converged to the optimal vocalization with respect to higher chance of response, whereas in the active sampling simulation, vocalizations oscillate around that perceived optimal vocalization with a set of quasi-optimal vocalizations. By adopting an active sampling strategy, the individual is maximizing learning by minimizing the uncertainty about the environment, since it is choosing the vocalization that minimizes the range of their beliefs about it. This strategy makes the marmoset vocal communication more robust in complex environments and thus more adaptive. This maximization of learning through vocalizations is also consistent with how human infants communicate with caregivers. Infant vocalizations often results in turn-taking with caregivers which provides a context for learning from those responses (38), and caregivers often change their responses to infant vocalizations to facilitate learning by the infant (39). As we are suggesting for marmoset vocal exchanges, these human infant-and-caregiver exchanges represent a feedback loop for learning about each other (40).

## Data, Materials and Software Availability

Data and analysis code to generate each of the figures is available at this link: https://github.com/ThiagoTVarella/Active-Sampling-in-Primate-Vocal-Interactions/tree/main

## Acknowledgments

We are very grateful to Gabriel Vercelli for help with the Bayesian modeling and to Steven Elmlinger for comments and suggestions on an earlier draft of this manuscript. This work was supported by an NSF Graduate Fellowship DGE-2039656 (TTV) and NIH-NINDS R01NS054898 (AAG).

## Methods

### 1. Dataset

The dataset reported here was reported previously (15). Recordings were obtained from 6 marmosets separated into 3 pairs of cagemates, each pair being one female and one male. The marmosets had ad libitum access to water and were fed with commercial marmoset diet, fresh fruits, vegetables, and insects. All experiments were performed with the approval of the Princeton University Institutional Animal Care and Use Committee.

### 2. Overview of the dyadic simulation algorithm

In this paper, we tested 3 different policies, which are 3 different ways to determine how to choose the duration of a vocalization given their previous knowledge. For this, we simulated 2 identical agents interacting with the same policy according to the following pseudocode:

**FOR** each time-iteration in the simulation:

**IF** agent 1 will vocalize:

**SET** probability distribution of the call duration given by policy and current belief

**SET** duration of vocalization based on the distribution

**STORE** duration of vocalization

**SET** whether agent 2 will respond

**UPDATE** belief of agent 1

**END IF**

**IF** agent 2 will vocalize:

**SET** probability distribution of the call duration given by policy and current belief

**SET** duration of vocalization based on the distribution

**STORE** duration of vocalization

**SET** whether agent 1 will respond

**UPDATE** belief of agent 2

**END IF**

**END FOR**

The details about the probability distribution are given in sections 4 of the methods for the active sensing policy, and section 5 for the vocal accommodation policy. The determination of whether the other agent will vocalize is explained in section 3 of the methods. The update of the previous knowledge is explained in section 6.

### 3. Sensing and observation model

We adapted the sensing model from Chen et al. (32). In our model, we assume that an agent is sensing the likelihood *V* to obtain a response from the other animal. After each vocalization, the marmoset receives a measurement (whether there was a response or not), which is used to inform a perceived response likelihood *V* to that vocalization.

To model *V*, let’s first define a function ϒ(*θ′*, x) that, given a sampled call duration *x* and the optimal call duration *θ′*, returns the assumed ground truth probability of getting a response to that vocalization. Considering that the measured response from the agent includes an error, we can define *V* =ϒ(*θ′*, x) + *ϵ*, where *ϵ* is a zero-mean Gaussian measurement noise. The function ϒ(*θ′*, x) can be assumed as the following Gaussian:

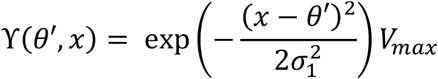

so that the minimum response probability is 0 when *x* is very different than *θ′*, and the response probability is maximum at *x* =*θ′*, which is defined as the parameter *V*_*max*_, given that no vocalization elicits a response 100% of the time. The parameter *σ*_1_ in the function is how broad that function is without changing the peak, which means that a big *σ*_1_ implies more values of*x* can be associated with a high response probability. In our simulations, we used *σ*_1_ =0.9, V_*max*_ =0.5. For our simulations, without loss of generality, the range of possible vocal durations is from 0 to 5, and the optimal duration is 1.5 (the values were chosen to be similar to real life values in seconds, even though the simulations did not assume a unit of measurement, nor is it intended to fit the exact values observed in real life). Note that the likelihood does not need to sum up to one as it is not a probability distribution.

### 4. Active sensing and the Expected Information Density (EID)

Consider that the marmoset has a belief about what that optimal vocal duration is, which is represented here by a probability distribution. If the marmoset only samples the duration where the belief is highest, small variations in the sampled vocalization will not lead to big differences in the measured response probability, because in the peak the belief is locally flat. If the vocalization sampled is in a region with higher slope, any small change on the sampled vocalization might lead to differences in the received information and less obvious results, which can imply more learning.

Following Chen et al. (32), we model the active sensing as a policy that maximizes the learning.

The belief, represented by *p*_*t*_(*θ*), is a probability distribution that informs the probability that each possible value *θ* is equal to the optimal duration *θ′*, at a time *t*. It starts as a uniform distribution, assuming that the agent doesn’t have any previous information about the physical and social environment. In real life, the marmoset generally will have some expectation about how to act where they are.We measure the uncertainty associated with a given belief *p*_*t*_ using the associated Shannon entropy, which is calculated via:

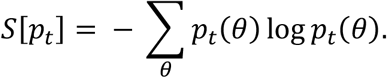

The lower the entropy is, the more certainty the agent has in the belief. We can describe the change in entropy as *ΔS*(*x*, V) =*S*[*p*_*t*_] − S[*p*_*t*+1_(*θ*|*V*, x)], where *p*_*t*+1_(*θ*|*V*, x) is the new belief of each possible value of *θ* after a given sampled *x* and the measured response likelihood *V*. This way, the best vocalization to maximize learning is the one that most lowers the expected entropy after the update, i.e., it lowers

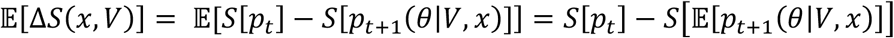

where 𝔼[*p*_*t+*1_(*θ*|*V*, x)] is the expected updated belief.

For each potential vocalization *x*, we iterate over possible values of *V* to calculate the expected updated belief given a possible *x* sampled and a hypothetical response likelihood *V*.

For a given *x* and *V, p*_*t+*1_(*θ*|*V*, x) is calculated applying a Bayesian filter to update *p*_*t*_(*θ*). The formula used is

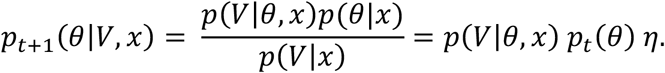

Here, *η* is just a normalization constant since the updated belief is still a probability distribution. We use that *p*(*θ*|*x*) =*p*_*t*_(*θ*), since *θ* does not depend on *x*. Finally, the function *p*(*V*|*θ*, x) is a likelihood function that informs how likely the marmoset is to actually associate the response probability *V* with the current belief of *θ*. Given that *V* =ϒ(*θ*, x) + *ϵ*, we can define *p*(*V*|*θ*, x) as a Gaussian likelihood centered in ϒ(*θ*, x) with a deviation *σ*_2_, so

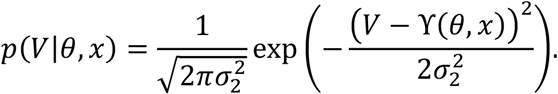

This allows us to calculate the entropy reduction *ΔS*(*x*, V) for each *x* and *V*. After calculating *p*_*t+*1_(*θ*|*V*, x) for each possible *V*, we calculate 𝔼[*p*_*t+*1_(*θ*|*V*, x)] by marginalizing over *V*. By integrating over possible values of *V*,we get a value for how much we expect each sampled vocalization to change the entropy, that is, *ΔS*(*x*), which we define as the *EID*(*x*). To do this, we first need a probability of *V* given *x*, p(*V*|*x*), which we marginalize over *θ* through the formula

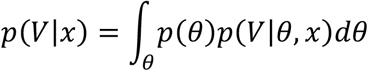

and we then use it as the weight of *ΔS*(*x*, V) in the integral that marginalizes over V in the interval from 0 to 1. In summary:

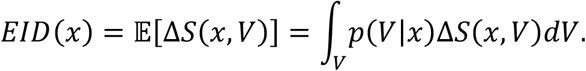

Only in the active sensing policy we calculate the *EID*(*x*). We do so for every *x* within the interval of 0 to 5, and we can define the result of this process as the distribution *EID*([0,5]). To make this calculation in our simulations, we used. σ_2_=0.3

The call duration is then sampled from the expected information density normalized to a probability distribution, i.e., *x* ∼ E*ID*([0,5]). That way, the vocalization is chosen from the interval [0,5] and is the one more likely to maximize the expected entropy reduction in the belief, i.e., the amount of new information that can be acquired from vocalizing within that interval.

### 5. Decision making for vocal accommodation

In all the policies, the call duration chosen for the next vocalization is sampled from a probability distribution. For the vocal accommodation policy, the call duration is sampled from the belief itself, i.e., *x* ∼ p_*t*_(*θ*). That way, vocalizations are chosen to maximize the expected response rate

### 6. Bayesian update of the belief

After the duration of the call is decided (*x*), we will observe whether the other agent produces a vocalization or not. For the active sensing and the vocal accommodation policies, the agent will update the belief depending on the response. We modeled the update using a Bayesian filter.

Let’s say that *r* ∈ Error! Bookmark not defined. is the variable is the variable that represents the response, *r* =1 if there is a response. To update the belief given the response, we can use the following Bayesian filter instead:

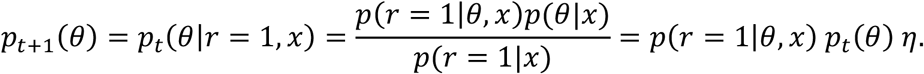

Notice however that *p*(*r* =1|*θ*, x) (the chance of receiving a response given *θ* and *x*) is directly related to the observational model ϒ(*θ*, x), so with *η* as a normalizing factor, we would have

*p*_*t*+1_(*θ*) =*η* ϒ(*θ*, x)*p*_*t*_(*θ*).

If there was no response, then:

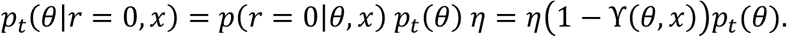

